# Human CST complex protects replication fork stability by directly blocking MRE11 degradation of nascent strand DNA

**DOI:** 10.1101/797647

**Authors:** Xinxing Lyu, Kai-Hang Lei, Olga Shiva, Megan Chastain, Peter Chi, Weihang Chai

## Abstract

Degradation and collapse of stalled replication forks are main sources of genome instability, yet the molecular mechanism for protecting forks from degradation/collapse is not well understood. Here, we report that human CST (CTC1-STN1-TEN1), a single-stranded DNA binding protein complex, localizes at stalled forks and protects forks from MRE11 nuclease degradation upon replication perturbation. CST deficiency causes nascent strand degradation, ssDNA accumulation after fork stalling, and delay in replication recovery, leading to cellular sensitivity to fork stalling agents. Purified CST binds to 5’ overhangs and directly blocks MRE11 degradation in vitro, and the DNA binding ability of CST is required for blocking MRE11-mediated nascent strand degradation. Finally, we uncover that CST and BRCA2 form non-overlapping foci upon fork stalling, and CST inactivation is synthetic with BRCA2 deficiency in inducing genome instability. Collectively, our findings identify CST as an important fork protector to preserve genome integrity under replication perturbation.

## Introduction

Faithful DNA replication is fundamental to genome integrity maintenance. During the process of DNA replication, replication machinery may encounter obstacles that perturb replication and cause replication forks slowing or stalling, leading to replication stress. Defects in stalled fork restart or stabilization induce genome instability, which is the main source of the pathology of human genetic diseases including cancer, aging, neurological diseases and developmental defects (Mazouzi et al, 2014; Zeman & Cimprich, 2013).

A panoply of mechanisms protects genome stability in response to fork stalling to guard normal cell proliferation. Slowing or stalled replication generates extensive ssDNA stretches at forks, which are bound by the replication protein A (RPA). RPA binding to ssDNA activates the ataxia telangiectasia and Rad3-related (ATR) kinase and recruits multiple proteins to facilitate the protection and restart of stalled forks (Flynn & Zou, 2011; Marechal & Zou, 2015). Multiple pathways, including dormant origin firing, replication repriming, translesion synthesis, template switching, and fork reversal, act to facilitate the protection and restart of the stalled forks (Bhat & Cortez, 2018; Zeman & Cimprich, 2013).

Electron microscopy analysis has revealed that various replication stressors can trigger fork remodeling to form a chicken foot-like physical structure through fork regression (Kolinjivadi et al, 2017; Zellweger et al, 2015). This fork reversal mechanism has emerged as an important means for stabilizing stalled forks and resuming replication following replication perturbation. For this reason, the dynamics of reversed forks has been extensively studied (Neelsen & Lopes, 2015; Quinet et al, 2017b; Rickman & Smogorzewska, 2019). The formation of reversed forks is catalyzed by SMARCAL1, ZRANB3, FANCM, HLTF, FBH1, and perhaps others (Betous et al, 2012; Gari et al, 2008; Kile et al, 2015; Kolinjivadi et al, 2017; Taglialatela et al, 2017; Vujanovic et al, 2017), and also involves RAD51 (Zellweger et al, 2015). While fork reversal is a protective mechanism to preserve fork stability under replication stress, the regressed arm on the reversal fork also provides a target for nuclease-dependent end resection. Without fork protectors such as BRCA1/2, RAD51, BARD1 and Abro1, reversed forks become vulnerable to nucleases MRE11, EXO1, DNA2, CtIP (Billing et al, 2018; Kolinjivadi et al, 2017; Lemacon et al, 2017; Mijic et al, 2017; Thangavel et al, 2015; Xu et al, 2017; Ying et al, 2012). Uncontrolled fork degradation by these nucleases is detrimental to genome stability and has been linked to both the lethality of BRCA2-defective embryonic stem cells and the sensitivity of BRCA-defective cells to certain chemotherapeutic treatments (Ray Chaudhuri et al, 2016; Schlacher et al, 2011; Ying et al, 2012). To date, the molecular mechanism underlying the balance between the protection and resection of regressed arms remains unclear.

The human CST complex, comprising of CTC1, STN1 and TEN1, contains multiple OB-fold domains and resembles the RPA70-RPA32-RPA14 complex structure (Miyake et al, 2009). CST prefers binding to G-rich sequences *in vitro* in a manner dependent on the sequence length (Chen et al, 2012; Miyake et al, 2009). It is capable of binding to an 18-nt G-rich ssDNA with high affinity, but the sequence specificity is lost if the oligonucleotide becomes longer (Hom & Wuttke, 2017; Miyake et al, 2009). Interestingly, although CST is unable to bind to dsDNA, ss-ds DNA junctions stabilize CST binding and decrease minimal ssDNA length requirement for CST binding to 10 nt (Bhattacharjee et al, 2017), indicating that CST may bind to special DNA structures with ss-ds junctions *in vivo*. A set of missense mutations of CTC1 and STN1 genes have been identified in patients with the Coats plus disease, a rare autosomal recessive disorder characterized by bilateral exudative retinopathy, retinal telangiectasias, growth retardation, intracranial calcifications, bone abnormalities, gastrointestinal vascular ectasias, accompanied by common early-aging pathological features (Anderson et al, 2012; Keller et al, 2012; Simon et al, 2016).

Like its homolog in budding yeast (Cdc13-Stn1-Ten1), the well-characterized and conserved roles of CST are in preserving telomere integrity. CST promotes efficient and complete replication of telomeric DNA to prevent sudden loss of telomeres (Gu & Chang, 2013; Huang et al, 2012). In addition, CST stimulates the priming activity of DNA polymerase α (POLα)-primase (Ganduri & Lue, 2017; Lue et al, 2014), and mediates C-strand fill-in at telomere ends to replenish resected C-strands (Huang et al, 2012). CST also inhibits telomerase access to telomeres and coordinates G- and C-strand synthesis (Chen et al, 2012).

We have previously shown that CST associates with G- or C-rich repetitive non-telomeric sequences in response to fork stalling, probably by binding to ssDNA accumulated at stalled forks (Chastain et al, 2016). CST facilitates fork recovery and its deficiency induces instabilities of these sequences and chromosome fragmentation, indicating that CST is an important player for protecting genome stability under replication stress (Chastain et al, 2016). However, the precise molecular mechanism underlying how CST participates in rescuing stalled replication remains unclear.

In this study, we report that CST is located at stalled forks upon hydroxyurea (HU) treatment, and CST deficiency leads to extensive ssDNA accumulation under replication stress and delays replication recovery. DNA fiber analysis demonstrates that CST depletion induces MRE11 degradation of nascent strand DNA that is dependent on fork reversal, suggesting that CST antagonizes nascent DNA degradation at reversed forks. Using purified human CST and human MRE11 proteins, we find that CST binds to the ss-ds DNA structure that mimics the reversed fork, directly blocking MRE11 degradation *in vitro*. The fork protection function of CST requires its binding to DNA, and loss of CST binding to DNA leads to nascent strand degradation in cells and *in vitro*. In addition, we uncover that CST inactivation exacerbates genome instability in BRCA2 deficient cells. Collectively, our results suggest that CST binds to reversed forks to inhibit MRE11 access to nascent strand DNA, thus acting as a direct fork protector to maintain genome stability in response to replication stress.

## Results

### CST localizes at stalled forks

We have previously shown that in addition to promoting efficient replication of telomeric DNA, CST is also enriched at G- or C-rich repetitive sequences genome-wide after replication fork stalling to protect the stability of these sequences (Chastain et al, 2016). However, it is unknown whether such protective function is direct or indirect, since there has been no direct evidence demonstrating that CST localizes at stalled forks. To gain insight into the function of CST in protecting genome stability, we examined CST localization at stalled sites in response to replication stress using the *in situ* protein interactions at nascent and stalled replication forks (SIRF) assay (Roy et al, 2018). SIRF offers sensitive visualization of protein localization at forks at a single-cell level if the protein-of-interest is in close proximity to EdU-labeled nascent strands. Briefly, nascent DNA in proliferating cells is labeled with EdU, and the incorporated EdU is then biotinylated by click reaction. Subsequently, the proximity ligation assay (PLA) is used to visualize the colocalization of target protein with biotinylated EdU (Fig. 1A). We constructed U2OS cells stably expressing Myc-CTC1 or Myc-STN1 via retroviral transduction, pulse-labeled cells with EdU for 8 min, treated them with HU for 3 hrs, and then carried out SIRF analysis using anti-Myc antibody (Fig. 1B). No signal was observed in the absence of EdU labelling or cells expressing vector only, suggesting that SIRF assay was specific for determining protein localization at replication forks (Fig. 1B). For the control, we performed SIRF against the DNA polymerase processivity factor proliferating cell nuclear antigen (PCNA). As expected, PCNA localized at active replication forks and formed distinct SIRF signals in untreated replicating cells (Figs. 1B and 1C). We also observed decreased PCNA SIRF foci after HU-induced fork stalling (Figs. 1B and 1C), consistent with previous findings from iPOND and SIRF assays that the amount of PCNA at stalled forks reduces when compared to that at elongating forks (Roy et al, 2018; Sirbu et al, 2011).

**Fig. 1.**
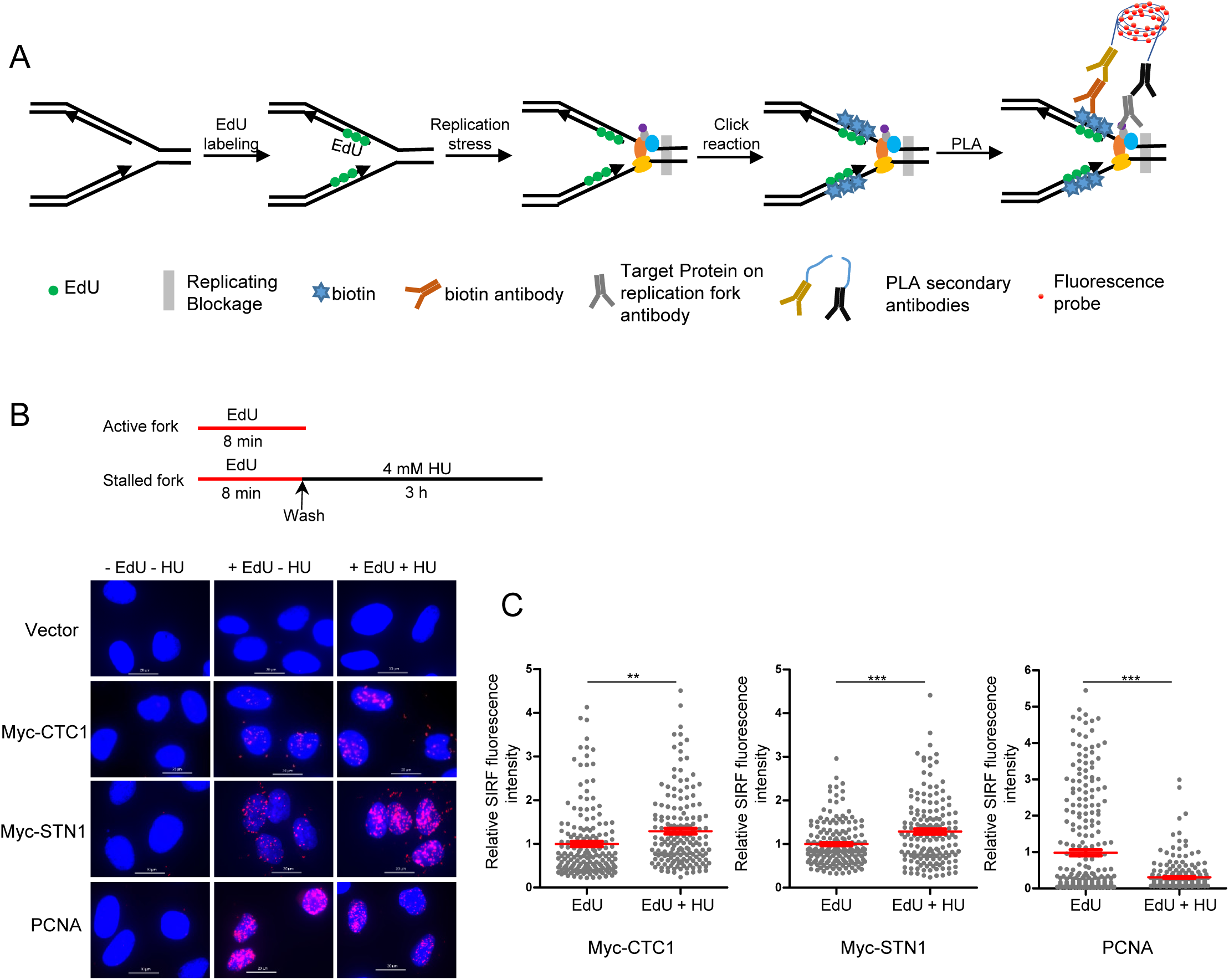
CST localizes at stalled forks. **(A)** Scheme of SIRF assay. Nascent DNA was pulse-labelled with EdU. Upon fork stalling, proteins located at stalled forks are in close proximity to EdU. EdU was then biotinylated using click reaction, followed by incubating with biotin and target protein antibodies. PLA amplification was used to visualize the localization of target proteins at EdU-labeled stalled forks. **(B)** Detection of CST at forks with SIRF assay. Cells stably expressing 13xMyc-CTC1 or 14xMyc-STN1 were used. To detect CST at stalled forks, cells were pulse-labeled with EdU for 8 min, followed by 4 mM HU treatment for 3 h. To detect CST at active forks, SIRF was performed without HU treatment. Representative images of vector only, 13Myc-CTC1, 14Myc-STN1 and PCNA on normal or stalled replication forks by SIRF analysis are shown. Scale bars: 20 µm. **(C)** Scatter plot of relative CTC1-SIRF, STN1-SIRF, and PCNA-SIRF fluorescence intensity with and without HU treatment. Relative SIRF fluorescence intensity was calculated by normalizing to respective untreated sample. In all SIRF experiments, three independent treatments were performed and results from one set of experiment are shown in each graph. P values were calculated by unpaired t-test with Welch’s correction. Red lines indicate the mean values. Error bars: SEM.

We detected SIRF signals of CTC1 and STN1 in untreated replicating cells, indicating that CST also localizes at normal unperturbed forks (Figs. 1B and 1C). This is consistent with previous reports that CST interacts with DNA polymerase α and facilitates DNA replication (Casteel et al, 2009; Chen et al, 2013; Ganduri & Lue, 2017). SIRF amplification signals from CTC1 and STN1 were increased by HU treatment (Figs. 1B and 1C), suggesting an increased CTC1 and STN1 localization at stalled forks in response to HU treatment.

### CST deficiency leads to the accumulation of ssDNA upon replication stress and results in delayed fork recovery

The initial outcome of replication stalling is the aberrant ssDNA production at stalled forks, likely caused by uncoupling of DNA helicase from DNA polymerase machinery. CST is a ssDNA binding protein complex and previously has been reported to suppress the formation of aberrant ssDNA at telomeres (Chen et al, 2013; Huang et al, 2012). We therefore examined the effect of CST depletion on ssDNA production after fork stalling in the genome. HeLa cells depleted of STN1 were treated with HU for 6 h, a condition that induced fork stalling and produced ssDNA (Ercilla et al, 2019) (Fig. 2A). Subsequently, BrdU staining was performed under the non-denaturing condition to measure ssDNA amount. As shown in Fig. 2B, CST depletion increased the amount of ssDNA after HU treatment, suggesting that CST loss leads to aberrant ssDNA accumulation (Fig. 2B).

**Fig. 2.**
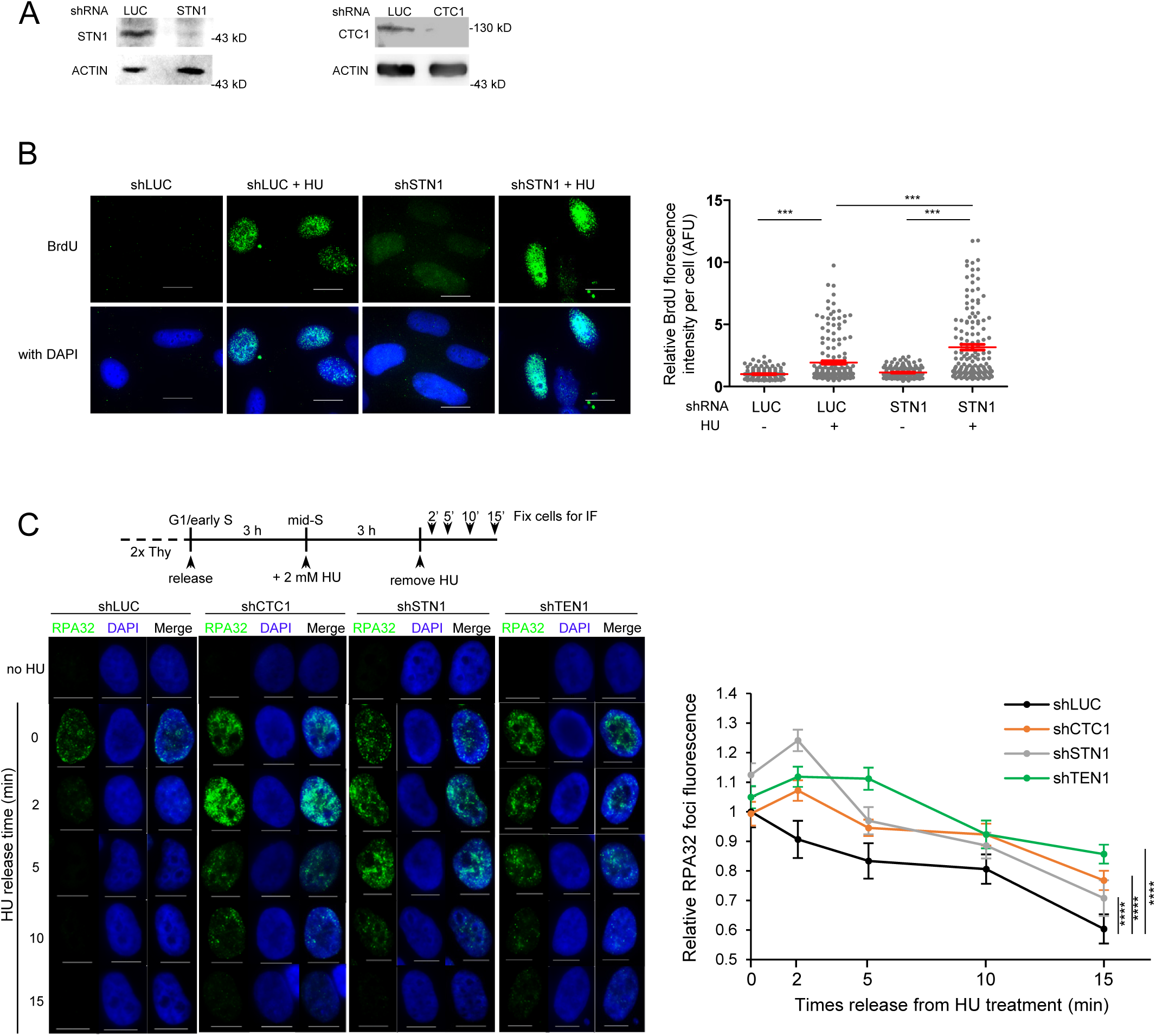
CST suppresses the accumulation of ssDNA at stalled forks. **(A)** Western blot showing STN1 and CTC1 knockdown. **(B)** CST deficiency causes ssDNA accumulation upon fork stalling. HeLa cells with STN1 knockdown were incubated with 10 µM BrdU for 48 h, followed by 2 mM HU treatment for 6 h, and subsequently stained with BrdU antibody under a non-denaturing condition to detect ssDNA. Scale bars: 20 µm. Amplified images showing distinct BrdU foci are included in Suppl. Figure S1. Relative BrdU fluorescence intensity was quantitated by Image J. *P* values were calculated by one-way ANOVA analysis with post hoc Tukey from three independent experiments. Error bars, SEM. **(C)** CST deficiency delays fork recovery. HeLa cells with CST knockdown were enriched in the mid-S phase as described in Methods, treated with 2 mM HU for 3 hrs to stall replication, and then released to no HU media for indicated time, fixed, and stained with RPA32 antibody. Scale bars in each image: 10 μm. Mean RPA32 nuclear foci fluorescence was quantified and relative foci fluorescence was calculated by normalizing to shLUC at 0 time point. Over all time points of *P* values were calculated by two-way ANOVA analysis with post hoc Tukey, p**** ≤ 0.0001. Error bars, SEM.

Since aberrant ssDNA accumulation upon fork stalling could impair fork recovery, we next examined whether CST depletion affected fork recovery by monitoring RPA32 foci disappearance after HU release. HeLa cells with CTC1, STN1, or TEN1 knockdown were first enriched in the early-S phase using double-thymidine as described previously (Chastain et al, 2016), and then released into the S phase for 3 h, at a time when the majority of cells reached the mid-S phase (Chastain et al, 2016). HU was added to media for an additional 3 h to stall replication. Following the removal of HU, cells were released into no-HU media to allow for fork recovery, and fixed at 2, 5, 10, 15 min immediately following release to detect RPA32 foci disappearance (Fig. 2C). While RPA32 foci quickly disappeared in control cells − indicative of rapid fork recovery, depletion of each subunit of CST significantly delayed RPA32 foci disappearance (Fig. 2C). Collectively, our results suggest that CST suppresses ssDNA accumulation and is important for preventing persistent fork stalling.

### CST antagonizes MRE11 degradation of nascent strand DNA at stalled forks

In response to replication stalling, forks may reverse to stabilize stalled forks and promote restart (Quinet et al, 2017b). Reversed forks, if not protected properly, are attacked by nucleases such as MRE11, DNA2 and EXOI, causing excessive degradation of nascent strand DNA (Lemacon et al, 2017; Thangavel et al, 2015). The accumulation of ssDNA and persistent RPA32 foci in CST deficiency cells led us to hypothesize that CST might play a role in protecting nascent strand DNA from nucleolytic degradation. To test this hypothesis, we analyzed fork stability using DNA fiber assays (Nieminuszczy et al, 2016; Quinet et al, 2017a). Replication tracks were sequentially labelled with chlorodeoxyuridine (CIdU) and iododeoxyurine (IdU) for 30 min, followed by 4 mM HU treatment for 5 h to induce fork stalling (Fig. 3A). Only bi-colored tracks were included in analysis. Using two independent shRNA sequences targeting STN1 (Fig. 3B), we found that STN1 depletion in a normal foreskin fibroblast cell line BJ resulted in the shortening of nascent strand DNA, and the presence of the MRE11 inhibitor mirin suppressed such shortening, indicating that STN1 depletion leads to MRE11-mediated degradation of nascent strand DNA at stalled forks (Fig. 3B). The same phenotype was recapitulated in two cancer cell lines, the human osteosarcoma cell line U2OS and colon cancer cell line HCT116 (Figs. 3C, 3D). Likewise, CTC1 depletion with shRNA caused similar fork degradation (Supplemental Fig. S2). Expression of the RNAi-resistant CTC1 (res-CTC1) cDNA fully rescued nascent strand shortening in knockdown cells, confirming that the phenotype was caused specifically by knockdown (Supplemental Fig. S2).

**Fig. 3.**
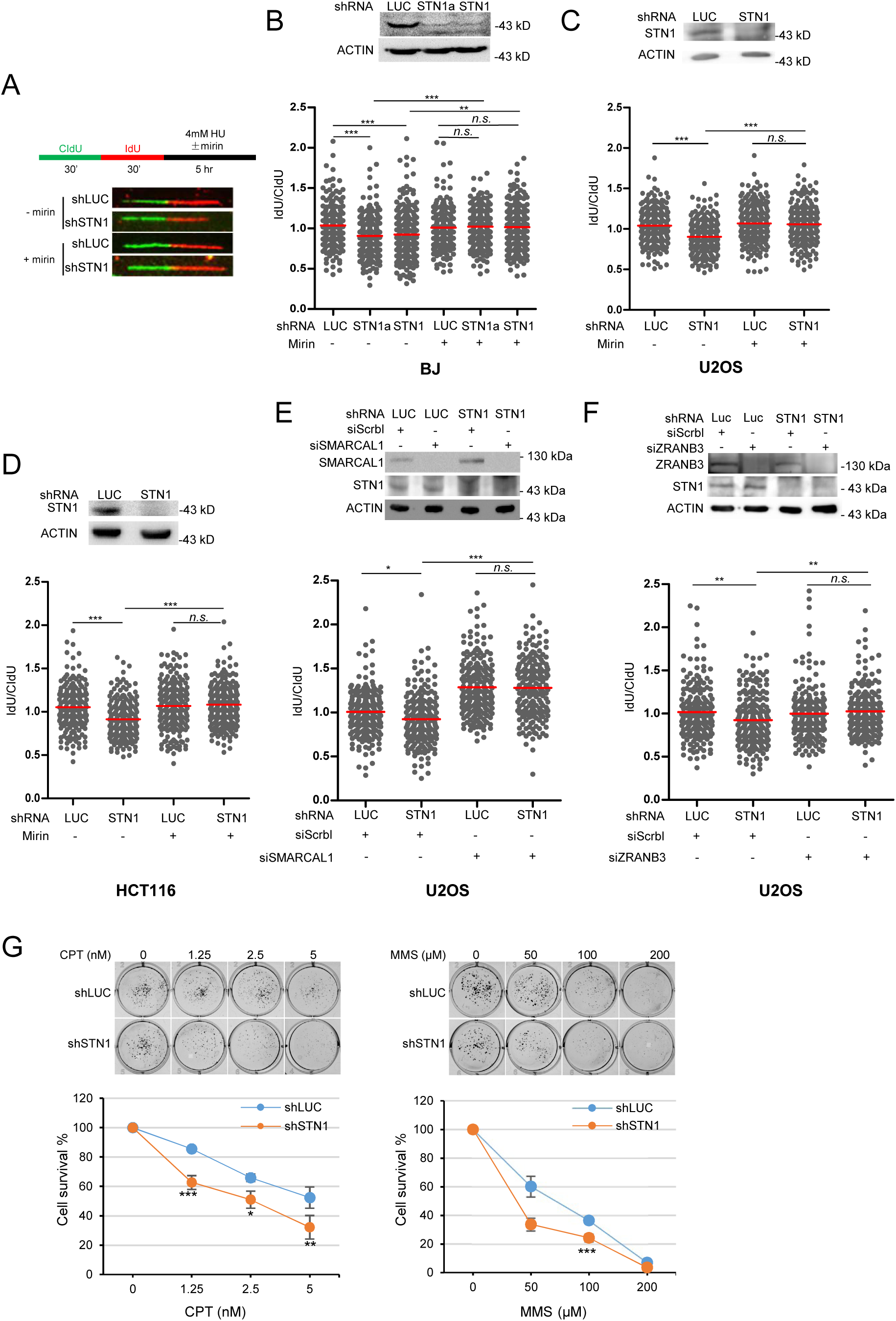
CST deficiency results in nascent strand degradation that is dependent on fork reversal, leading to sensitivity to fork stalling reagents. **(A)** Scheme of DNA fiber analysis and representative fiber images. **(B)** DNA fiber analysis of BJ cells with STN1 knockdown with and without mirin treatment. In all DNA fiber experiments in this study, at least two independent treatments and fiber experiments were performed to ensure reproducibility. Results from one set of experiment are shown in each graph. Scatter plots indicates IdU/CIdU tract length ratios for individual replication forks. *P* values were calculated by one-way ANOVA analysis with post hoc Tukey. Red lines indicate the mean values. Western blots showing protein knockdown in DNA fiber assays are included. **(C)** DNA fiber analysis of U2OS cells with STN1 knockdown with and without mirin treatment. **(D)** DNA fiber analysis of HCT116 cells with STN1 knockdown with and without mirin treatment. **(E)** DNA fiber analysis of U2OS cells showing that SMARCAL1-mediated fork reversal is needed for nascent stand degradation observed in CST-deficient cells. **(F)** DNA fiber analysis of U2OS cells showing that ZRANB3-mediated fork reversal is needed for nascent strand degradation observed in CST-deficient cells. **(G)** Colony formation assay showing that STN1 depletion decreases cell viability in U2OS cells after exposure to CPT and MMS. Results are the average of three independent experiments, with triplicate in each experiment. *P* values were calculated by one-way ANOVA analysis with post hoc Tukey from three independent experiments. Error bars, SEM.

### Nascent strand degradation caused by CST deficiency requires fork reversal and CST deficiency sensitizes cells to fork stalling agents

Fork reversal through remodeling of stalled forks into four-way junction structures is catalyzed by SMARCAL1, ZRANB3 and others (Kolinjivadi et al, 2017; Taglialatela et al, 2017; Vujanovic et al, 2017). To determine whether the fork protection function of CST relied on fork reversal, we knocked down SMARCAL1 or ZRANB3 in STN1-depleted cells and measured nascent strand degradation. Our results revealed that depletion of SMARCAL1 or ZRANB3 rescued nascent strand degradation in STN1-deficient cells (Figs. 3E, 3F). Consistent with previous reports (Coquel et al, 2018; Tonzi & Yin, 2018), depletion of SMARCAL1 slightly increased IdU/CldU ratio in control knockdown cells (Fig. 3E), though the underlying reason is unclear. Regardless, our results suggest that fork reversal is a pre-requisite for fork degradation in CST-deficient cells.

Stalled forks are prone to induce DNA lesions that are detrimental to genome integrity. Therefore, defects in replication stress response proteins are more sensitive to genotoxins. We found that STN1 knockdown cells displayed sensitivity to fork-stalling agents camptothecin (CPT) and methyl methanesulfonate (MMS), detected by the colony formation assay (Fig. 3G). Together, our data suggest that CST protects reversed forks from MRE11 nuclease degradation to maintain genome stability.

### The DNA binding ability of CST is required for protecting nascent strand degradation

CST is a ssDNA binding protein complex containing multiple OB folds (Bryan et al, 2013), and its DNA binding ability primarily relies on the two predicted OB-fold domains at the N-terminus of CTC1 (Miyake et al, 2009). To test the role of CST binding to ssDNA in antagonizing MRE11 degradation of nascent strand DNA, we constructed a truncated CTC1 mutant that removed the 700 residues in the N-terminus harboring the DNA binding domain (Fig. 4A). Since CST binds to telomeric DNA, we first tested whether Δ700N localized at telomeres following the expression of Myc-Δ700N in U2OS cells using retroviral transduction. As shown in Fig. 4B, while the full-length WT-CTC1 displayed telomere localization in both untreated and HU-treated cells, Myc-Δ700N exhibited diffused staining and failed to colocalize with TRF2, indicating that Δ700N failed to bind to telomeres (Fig. 4B). Moreover, purified Δ700N protein completely lost the DNA binding ability *in vitro*, validating that this mutant does not bind to DNA (Fig. 5D, see below). Next, we used co-IP to confirm that Δ700N was still able to form a complex with STN1 and TEN1. Myc-Δ700N was co-transfected with His_6_-STN1 and HA-TEN1 into HEK293T cells, followed by co-IP. As shown in Fig. 4C, Δ700N interacted with STN1 and TEN1 just like WT-CTC1. Thus, the CST complex formation was unaffected after deleting the DNA-binding domain of CTC1. Since CST interacts with RAD51 in response to HU treatment and this interaction is important for recruiting RAD51 to fragile sites (Chastain et al, 2016), we also tested the effect of Δ700N on RAD51 binding. Previously we have shown that deleting the N-terminal 600 amino acids of CTC1 has no effect on CST-RAD51 interaction while removing the N-terminal 840 amino acids partially loses RAD51 interaction (Wang & Chai, 2018). When Δ700N was tested, we found that Δ700N largely retained the RAD51 interaction (Fig. 4D), consistent with our previous finding that the RAD51-interacting domain primarily resides within the C-terminal half of CTC1 (Wang & Chai, 2018). This result also confirms our previous observation that CST interacts with RAD51 in a DNA-independent manner (Chastain et al, 2016).

**Fig. 4.**
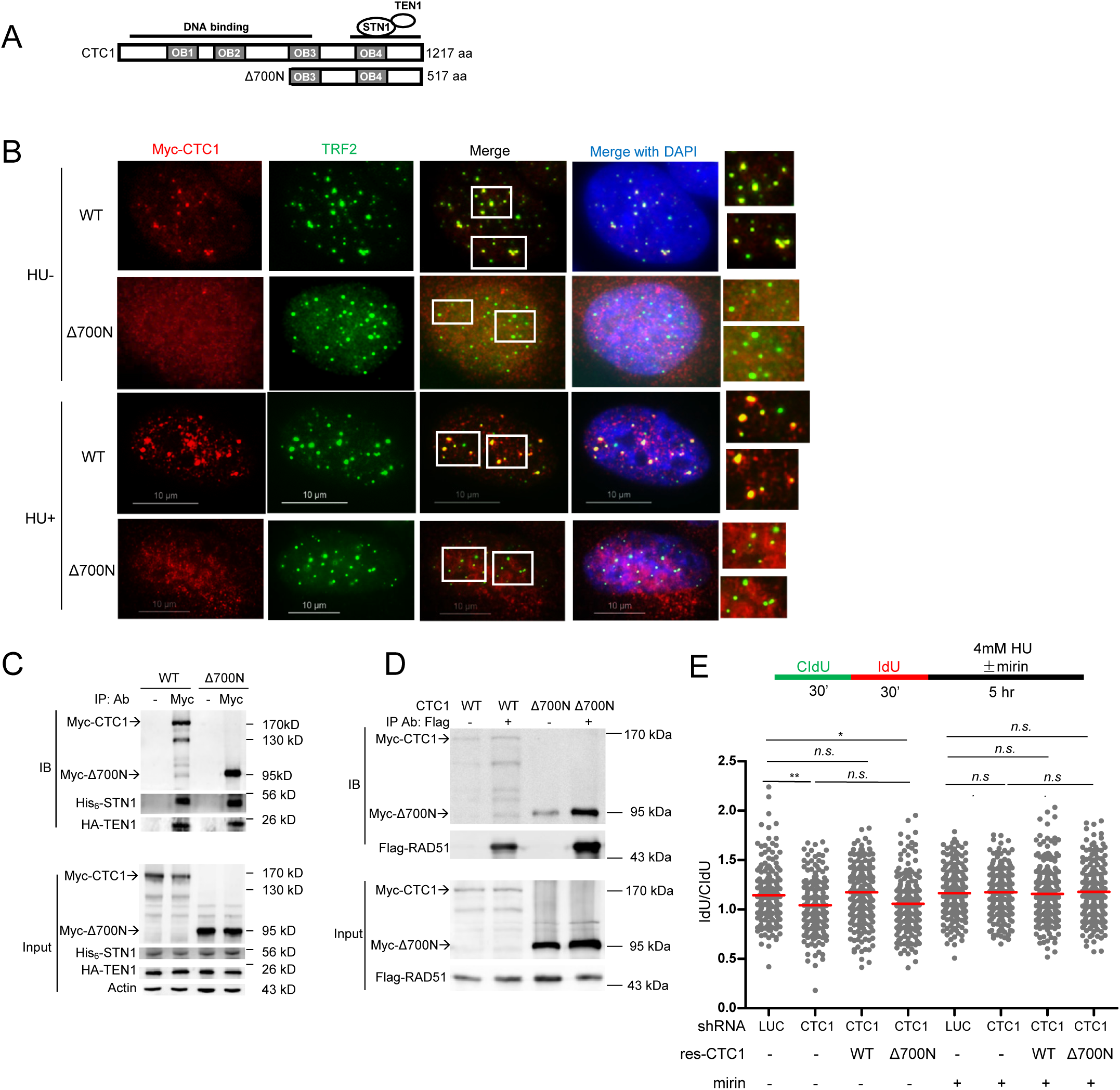
The ssDNA binding ability of CST is required for protecting nascent strand from degradation. **(A)** Illustration of the full-length CTC1 gene and Δ700N. STN1/TEN1 interacts with CTC1 via the predicted OB4 domain at the C-terminus, whereas the predicted N-terminal OB1 and OB2 domains are needed for DNA binding. **(B)** Deletion of the N-terminal 700 aa of CTC1 abolishes CTC1 localization at telomeres. U2OS stably expressing WT Myc-CTC1 and Myc-Δ700N were treated with or without HU and co-stained with Myc (red) and TRF2 (green) antibodies. Boxed areas are amplified in inserts to indicate CTC1/TRF2 colocalization (yellow). Scale bars: 10 μm. **(C)** Δ700N does not affect CST complex formation. HEK293T cells were co-transfected with Myc-CTC1, His_6_-STN1 and HA-TEN1. Co-IP was performed with Myc antibody to pulldown His_6_-STN1 and HA-TEN1. Full-length Myc-CTC1 was prone to degradation during immunoprecipitation. **(D)** Δ700N retains RAD51 interaction. HEK293T cells were co-transfected with Myc-CTC1, His_6_-STN1, HA-TEN1 and Flag-RAD51, and treated with HU (2 mM, 16 h). Co-IP was performed with Flag antibody to pulldown Myc-CTC1. **(E)** DNA fiber analysis of U2OS CTC1 knockdown cells rescued by RNAi-resistant WT and Δ700N. *P* values were calculated by one-way ANOVA analysis with post hoc Tukey. Red lines indicate the mean values.

**Fig. 5.**
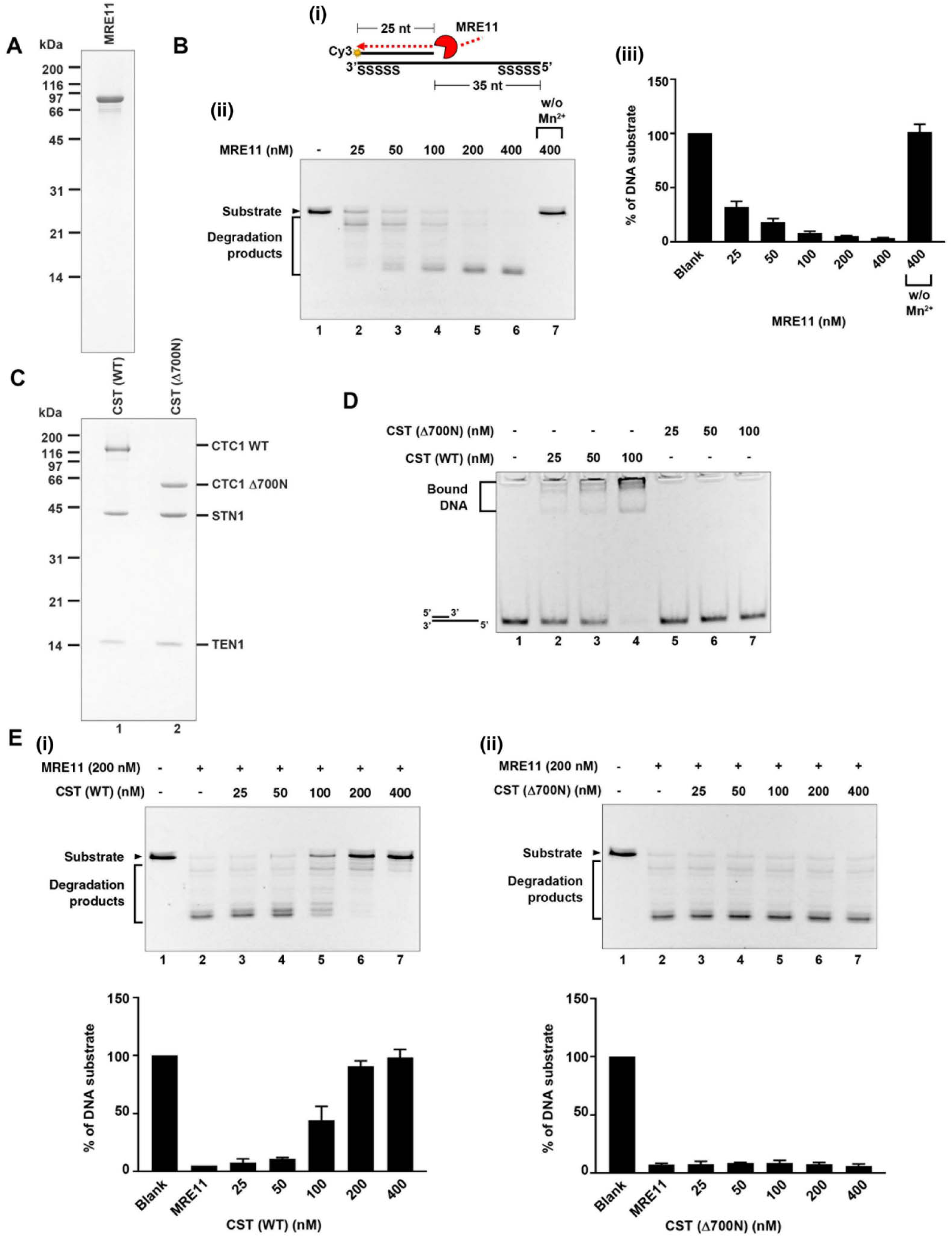
CST binding to 5’ overhangs directly blocks MRE11 degradation. **(A)** Coomassie blue stained SDS-PAGE gel (15%) of purified human MRE11. **(B)** (i) Scheme shows the nuclease activity of MRE11 in degrading 5’ Cy3-labeled substrates (25 nt + 60 nt with phosphorothioate bonds on both ends). (ii) The 5’ Cy3-labeled substrates (80 nM) were incubated with the indicated concentration of MRE11 with or without the Mn^2+^ cofactor at 37°C for 20 min. Reactions were stopped by SDS and proteinase K and resolved on 27% denatured polyacrylamide gel. (iii) The results are graphed, and error bars represent the standard deviation (± SD) calculated from at least three independent experiments. **(C)** Coomassie blue stained SDS-PAGE gel (15%) of purified human CST complex (CTC1 wild-type and Δ700N). **(D)** DNA binding activity of wild-type CST complex and Δ700N mutant complex. The 5’ Cy3-labeled substrates (80 nM) were incubated with the indicated concentrations of CST. Samples were analyzed by 6% native polyacrylamide gel. **(E)** MRE11 degradation analysis. 5’ Cy3-labeled substrates (80 nM) were pre-incubated with indicated concentrations of CST wild-type (i) or Δ700N mutant variant (ii) at 37°C for 5 min. Reactions were completed by adding MRE11 (200 nM) for an additional 20 min incubation and then stopped by SDS and proteinase K. Samples were resolved in 27% denatured polyacrylamide gel. The results are graphed and error bars represent the standard deviation (± SD) calculated from at least three independent experiments.

Next, we tested the effect of loss of CST binding to DNA on nascent strand degradation. RNAi-resistant WT-CTC1 or Δ700N were stably expressed in U2OS cells via retroviral transduction, and endogenous CTC1 was then depleted with shRNA. DNA fiber analysis showed that, while WT-CTC1 fully rescued nascent strand degradation caused by CTC1 knockdown, expressing Δ700N failed to rescue (Fig. 4E). Thus, our results suggest that CST binding to ssDNA is required for protecting nascent strand from MRE11 attack.

### Purified CST binds to 5’ overhang and directly blocks MRE11 degradation of DNA

The above observations indicated that CST binding to DNA could protect reversal forks from MRE11 degradation to maintain genome stability. MRE11 possesses the 3’ to 5’ exonuclease activity and can extensively degrade ss-ds DNA junction structures with 5’ overhangs formed by regressed forks (Kolinjivadi et al, 2017). It has been reported that CST prefers binding to ss-ds DNA junctions *in vitro* (Bhattacharjee et al, 2017). Hence, we hypothesized that CST might bind to 5’ overhangs of the regressed arms to protect nascent strand DNA from MRE11 degradation. To test this, we purified human MRE11 protein (Fig. 5A), generated a ss-ds DNA junction structure with a 5’ overhang that mimicked the regressed arm (Fig. 5B i), and used the purified human heterotrimeric CST complex and MRE11 proteins to reconstitute nascent strand DNA protection *in vitro*. Recombinant human MRE11 was purified from Expi293F cells as described in Methods (Fig. 5A), and then incubated with the ss-ds DNA substrate. In agreement with the previous report, MRE11-dependent degradation in the 3’ to 5’ direction was observed (Fig. 5B ii), and the MRE11 nuclease activity required cofactor Mn^2+^ (Fig. 5B iii) (Paull & Gellert, 1998). Next, we purified the human CST complex, both the WT-CST and Δ700N-ST from Expi293F cells to near homogeneity (Fig. 5C). To ensure that only the complete protein complexes containing all CST subunits were used in the biochemical assays, affinity-purified CST proteins were subject to gel filtration chromatography at the final step to remove non-complexed individual proteins. As shown in Fig. 5C, Δ700N was able to form the complex with STN1 and TEN1, validating our co-IP finding that Δ700N did not disrupt CST complex formation (Fig. 4C). While purified WT-CST complex bound to the 5’ overhang structure, Δ700N-ST failed to bind (Fig. 5D). Intriguingly, when incubating WT-CST pre-assembled ss-ds DNA junctions with MRE11, WT-CST protected DNA from MRE11 degradation in a dosage-dependent manner (Fig. 5E i). In contrast, Δ700N-ST completely failed to do so (Fig. 5E ii). These results support that CST binding to 5’ overhangs directly protects ssDNA from MRE11 degradation.

### Genetic relationship between CST and BRCA2 in maintaining genome stability

BRCA2 is another ssDNA binding protein which antagonizes nascent strand degradation (Schlacher et al, 2011). Interestingly, CST shares a number of molecular features with BRCA2: they both interact with RAD51, bind to ssDNA, assist RAD51 recruitment to stalled forks (Schlacher et al, 2011; Taglialatela et al, 2017). Additionally, the phenotypes found after CST knockdown at stalled forks are reminiscent to BRCA2-deficient cells. These observations prompted us to explore their genetic relationship in genome maintenance. We first analyzed genome instability in cells co-depleted of STN1 and BRCA2. As expected, depletion of STN1 alone or BRCA2 alone increased genome instability markers including micronuclei and abnormal anaphase bridges (Figs. 6A, 6B). Single depletion of either STN1 or BRCA2 also induced the accumulation of γH2AX-positive cells (Fig. 6C), indicative of increased spontaneous DNA damage that might have arisen from fork instability. When BRCA2 and STN1 were co-depleted, we observed elevated levels of genome instabilities measured by micronuclei, anaphase bridges, and γH2AX (Figs. 6A-6C). Collectively, these results suggest a synthetic effect of CST inactivation and BRCA2 deficiency in inducing genome instability. In addition, co-depletion significantly reduced BrdU incorporation (Fig. 6D), suggesting that CST and BRCA2 double-deficiency severely impairs DNA synthesis and diminishes the population of replicating cells. Despite our repeated attempts to use DNA fiber analysis to determine nascent strand degradation in double knockdown cells, we were unable to obtain data that truly represented fork dynamics due to the low number of replicating cells following double knockdown.

**Fig. 6.**
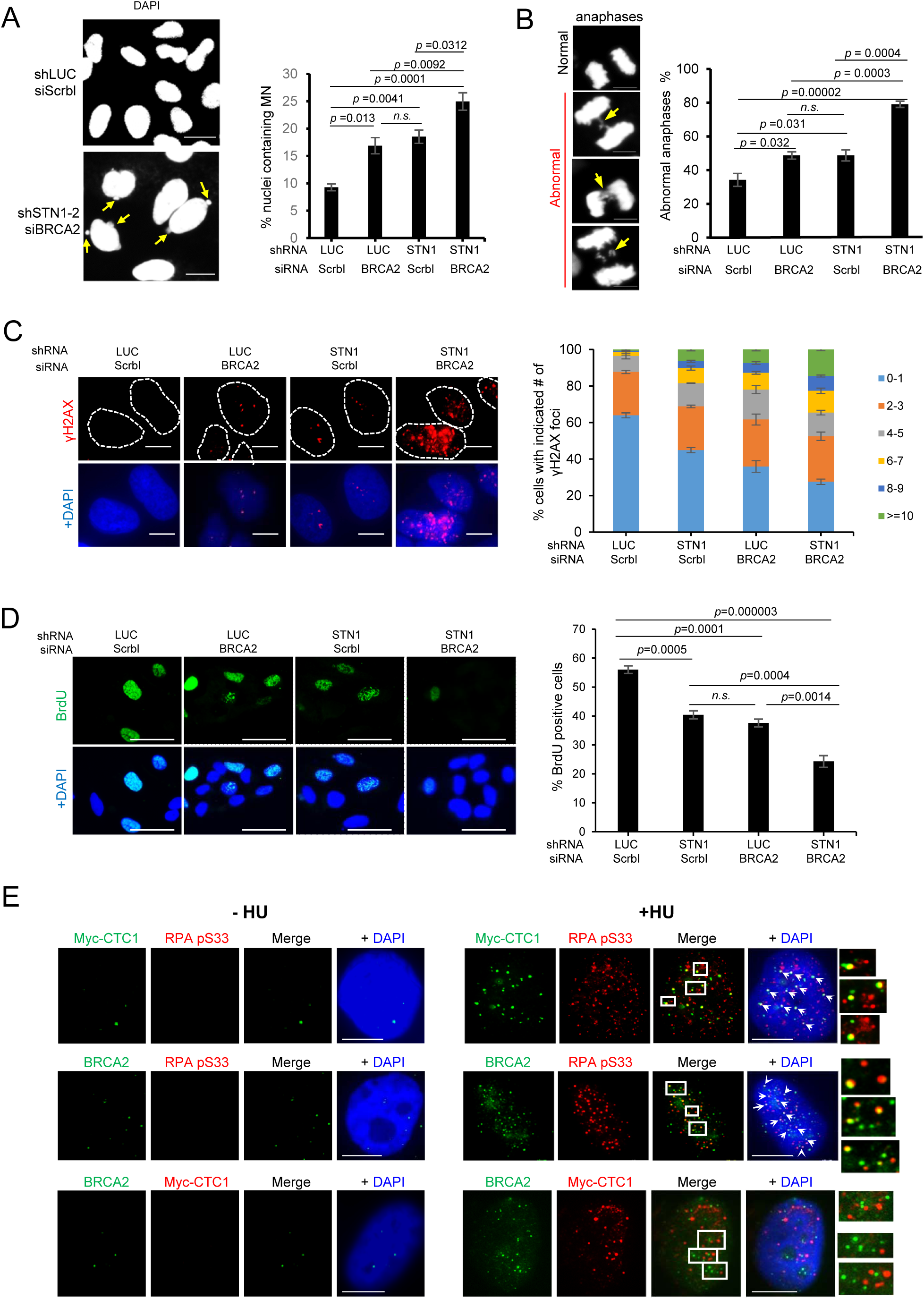
Genetic and spatial relationship of CST and BRCA2 in protecting genome stability. **(A)** Micronuclei (MN) formation in BRCA2- and STN1-deficient cells. BRCA2 and STN1 were co-depleted in U2OS cells with siRNA or shRNA. Representative DAPI staining images are shown. Arrows point to MN. Scale bars: 20 µm. Average percentage of nuclei containing MN are shown. *P* values were calculated by one-way ANOVA analysis with post hoc Tukey from three independent experiments. Error bars: SEM. **(B)** Anaphase bridges (arrows) in BRCA2- and STN1-deficient U2OS cells. Results were from three independent knockdown experiments. Scale bar: 10 µm. *P* values were calculated by one-way ANOVA analysis with post hoc Tukey from three independent experiments. Error bars: SEM. **(C)** γH2AX induced by BRCA2 knockdown and STN1 knockdown in U2OS cells. Results were from three independent knockdown experiments. *P* values were calculated by one-way ANOVA analysis with post hoc Tukey from three independent experiments. Error bars: SEM. **(D)** Co-depletion of STN1 and BRCA2 significantly impairs DNA replication. Results were from three independent knockdown experiments. Scale bar 50 µm. *P* values were calculated by one-way ANOVA analysis with post hoc Tukey from three independent experiments. Error bars: SEM. **(E)** BRCA2 and CST form non-overlapping foci after HU treatment. U2OS cells stably expressing Myc-CTC1 were treated with or without HU (2 mM) and stained with anti-Myc, anti-BRCA2, and anti-pS33. Boxed areas are amplified and shown at the right side of images. White arrows point to colocalizations. Scale bars: 10 µm.

### Spatial relationship between CST and BRCA2

It has been shown that CST prefers binding to DNA with G-rich repetitive sequences and structural features including ss-ds junctions (Bhattacharjee et al, 2017; Chastain et al, 2016; Hom & Wuttke, 2017). We postulated that CST might localize at forks stalled at non-BRCA2 binding regions and be particularly important for protecting the stabilities of these sequences. If so, CST and BRCA2 would be expected to localize at distinct loci after fork stalling. We thus used IF to test the spatial relationship of CST and BRCA2 upon fork stalling. Cells expressing Myc-CTC1 were treated with HU and then co-stained with BRCA2, Myc-CTC1, and S33-phosphorylated RPA32 (pS33). pS33 was used because it is detectable at stalled forks by iPOND within 10 min after HU addition (Sirbu et al, 2011), colocalizes with the BrdU-labeled ssDNA induced by HU treatment (Supplemental Fig. S3), and therefore marks ssDNA formed at stalled forks (Vassin et al, 2009). IF results show that after HU treatment, both BRCA2 and CST formed foci that colocalized with pS33 (Fig. 6E). This result further confirms our SIRF analysis finding that CST localizes at stalled forks (Fig. 1). Notably, CST or BRCA2 colocalized with a subset of pS33, and meanwhile their foci were largely non-overlapping (Fig. 6E). Our results suggest a mutually exclusive physical relationship between CST and BRCA2: they either localize and protect stalled forks at spatially distinct regions in the genome, or they may be independently recruited to the same site following fork stalling but at different stages.

### CST expression level positively correlates with cancer patient survival

Since genome instability gives rise to mutations that drive tumor initiation and development, we then assessed whether increased genome instabilities caused by CST deficiency would impact the outcome of cancer patient survival. Using Kaplan-Meier analysis, we found that breast cancer patients with CTC1 and STN1 expression at the upper tertile showed significant better disease-free survival, while a worse survival outcome was observed in patients with CTC1 and STN1 expression at the lower tertile (Supplemental Fig. S4). Future studies to determine whether CST can be used as a predictive biomarker in these subgroups may be clinically valuable.

## Discussion

Protecting stalled forks from excessive nucleolytic resection is critical in resolving replication stress and maintaining genome stability. A number of studies have shown that many homology-directed repair (HDR) factors, including BRCA1/2, RAD51, CtIP, have HDR-independent roles in protecting fork stability (Hashimoto et al, 2010; Kolinjivadi et al, 2017; Przetocka et al, 2018; Taglialatela et al, 2017). In this study, we provide direct evidence that the ssDNA-binding complex CST is a new component at stalled forks and functions in limiting excessive nascent strand degradation. We find that CST localizes at stalled replication forks. CST deficiency results in nascent strand degradation at reversed forks, leading to persistent ssDNA accumulation and delay in replication recovery. Consequently, CST depletion sensitizes cells to fork stalling agents. Using purified CST proteins and a reconstituted biochemical system, we also show that CST binds to 5’ overhangs and directly blocks MRE11 nucleolytic degradation of DNA *in vitro*, supporting our DNA fiber results. Collectively, results in this study suggest that CST binds to 5’ overhangs formed at reversed forks, and such binding directly blocks MRE11 and perhaps other nucleases from excessively resecting nascent strand DNA (Fig. 7).

**Fig. 7.**
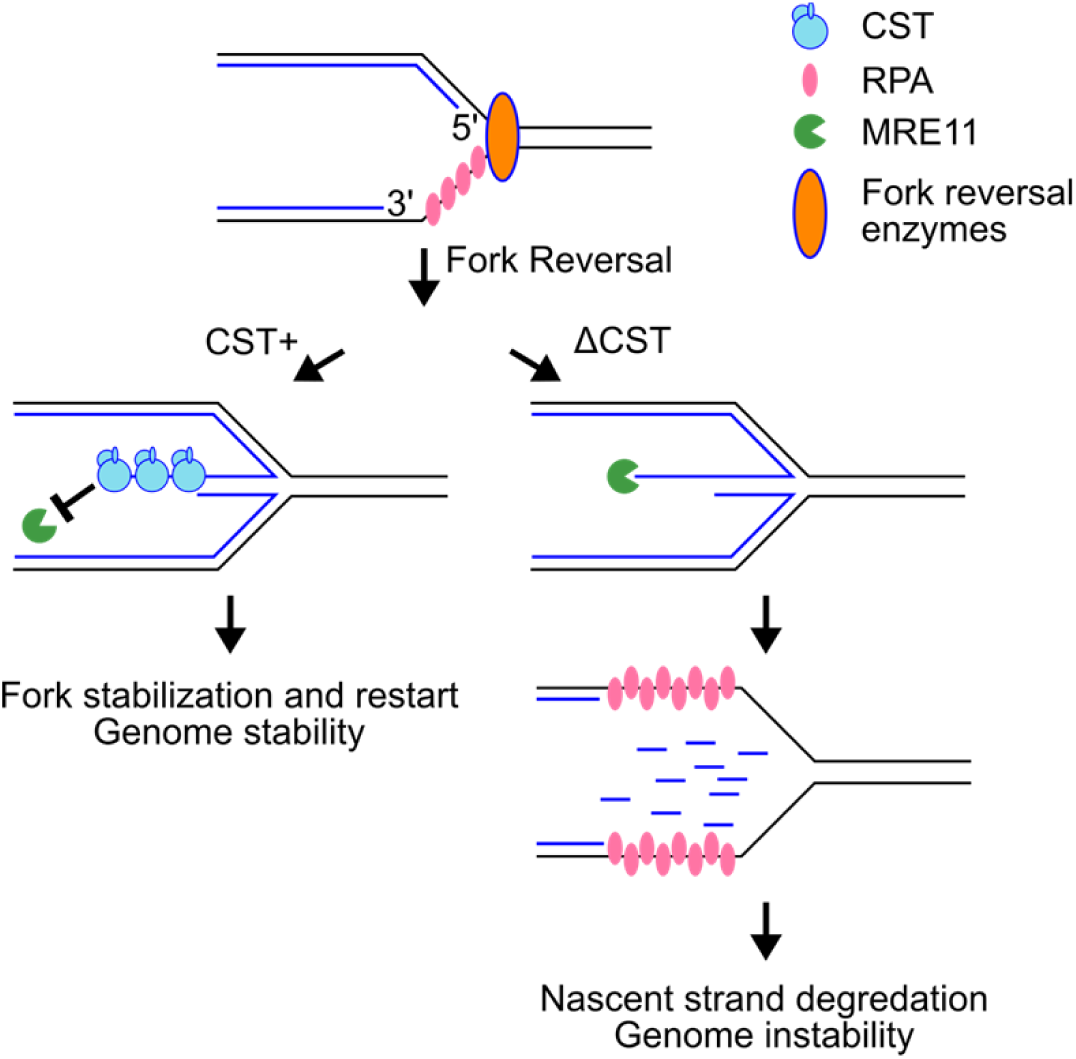
Model for CST in protecting reversal forks stability. Upon fork stalling, the stalled fork is reversed by SMARCAL1, ZRANB3, or others. CST binds to ssDNA (5’ overhang) at reversed forks to prevent MRE11 excessive degradation of nascent DNA strand. Without CST, nascent strand DNA is degraded by MRE11, leading to the generation of excessive ssDNA and genome instability.

CST was identified as an RPA-like protein complex due to its ssDNA binding ability and structural similarity to RPA (Miyake et al, 2009). Despite this similarity, recent discoveries on CST at stalled forks and DSBs suggest that CST is functionally distinct from RPA ((Barazas et al, 2018; Chastain et al, 2016; Mirman et al, 2018) and this study). RPA has no obvious ability to antagonize strand degradation (Kolinjivadi et al, 2017), nor does it play a role in inhibiting DSB end resection. In fact, RPA has been found to promote end resection (Cannavo et al, 2013; Yan et al, 2011). Although it has been well established that the primary function of RPA binding to DNA is for activating ATR (Marechal & Zou, 2015), there has been no report describing the participation of CST in ATR signaling. While the exact relationship of RPA and CST remains to be determined, we notice that CST shares a number of functional and structural similarities with another genome stability protector, BRCA2. Structurally, both BRCA2 and CST contain multiple OB-fold domains. Biochemically, they both bind to ssDNA and also interact with RAD51 (Chastain et al, 2016; Schlacher et al, 2011). Functionally, they both assist RAD51 loading to stalled sites and protect nascent strand DNA from MRE11 nucleolytic degradation, and CST depletion induces similar genome instabilities like BRCA2 depletion ((Schlacher et al, 2011) and this study). Interestingly, when we analyze CST and BRCA2 localization after fork stalling, we find that CST and BRCA2 localize at spatially distinct regions after fork stalling. Additionally, CST inactivation further elevates genome instability and severely impairs DNA replication in BRCA2-deficient cells (Fig. 6). Given that CST prefers binding to DNA with specific sequence (G-rich repetitive) and structural features (ss-ds junctions) (Bhattacharjee et al, 2017; Chastain et al, 2016; Hom & Wuttke, 2017) while BRCA2 binds to G-rich oligos with low affinity (Yang et al, 2002), it is possible that CST may be a surrogate of BRCA2 that is particularly important for maintaining the stability of G-rich repetitive sequences and/or sequences prone to form special structures – an interesting hypothesis to be further tested. Alternatively, it remains a possibility that CST and BRCA2 may occupy the same stalled site at different time following fork stalling.

CST interacts with POLα and stimulates POLα priming activity and primase-to-polymerase switching (Casteel et al, 2009; Ganduri & Lue, 2017; Lue et al, 2014; Nakaoka et al, 2011). At telomere ends, both CST and POLα participate in filling in C-strands at resected telomere ends to shorten telomeric G-overhang lengths (Gu et al, 2012; Huang et al, 2012). At DSBs in BRCA1-null cells, CST deficiency results in an increase of phosphorylation of RPA S4/S8, indicative of extensive end resection (Mirman et al, 2018). The resemblance of lengthened G-overhangs to increased DSB end resection after CST loss has led to the proposal that CST may also promote the fill-in synthesis at resected DSB ends, and that the defective fill-in caused by CST deficiency may give rise to long 3’ overhangs that are suitable for initiating strand invasion and HDR (Barazas et al, 2018; Mirman et al, 2018). Since the same group of nucleases involved in DSB end processing, namely MRE11, DNA2, and EXO1, also resect reversed forks, it is tempting to speculate that CST may stimulate POLα fill-in activity at reversed forks to antagonize end resection. Interestingly, our results show that CST binding to reversed forks can directly block MRE11 degradation of DNA (Fig. 5). While our present data do not exclude the possible involvement of fill-in synthesis in countering fork degradation, our results uncover a new function of CST in protecting fork stability and offer an alternative model that CST binds to ssDNA at reversed forks and directly inhibits MRE11 nuclease degradation, thereby protecting fork stability in a POLα-independent manner (Fig. 7). We speculate that CST occupying 5’ overhangs at reversed forks may establish a physical barrier for MRE11 access to the 3’ end. Future structural studies on the CST-DNA complex will be helpful to determine the exact nature of such a barrier.

Mutations of CST have been identified in the rare genetic disease Coats plus, and thus CST has been mostly studied in the context of Coats plus pathogenesis. Our findings identifying CST’s role in protecting the fork stability thus reaffirm CST as an important genome maintenance player. Since the instability of stalled forks is a major source of genome instability in the early stages of tumorigenesis, it is likely that CST dysfunction may contribute to carcinogenesis. Indeed, the Cancer Genomics Atlas database shows that the CST genes are frequently altered in many types of cancers with mutations, deletions and amplifications (Supplemental Fig. S5) (Cerami et al, 2012; Gao et al, 2013). The positive correlation between CST gene expression level and the survival outcome in breast cancer patients (Supplemental Fig. S4) suggests that CST may be worth clinical investigation as a potential prognosis biomarker. Investigating whether CST deficiency may promote tumorigenesis *in vivo* will also be compelling and crucial.

## Materials and Methods

### Cell culture

HeLa, U2OS, HCT116, BJ, HEK293T cells were obtained from American Type Culture Collection. Cells were cultured in DMEM media supplemented with 10% cosmic calf serum (ThermoFisher) at 37 °C containing 5% CO_2_.

### Plasmids

RNAi-resistant pcDNA-Flag-CTC1, pCL-CTC1-13xMyc, pcDNA-HA-TEN1, pBabe-Flag-STN1, pBabe-HA-TEN1 were described previously (Chastain et al, 2016; Huang et al, 2012; Miyake et al, 2009; Wang & Chai, 2018). The pCL-CTC1Δ700N-13xMyc plasmid was constructed by PCR amplifying CTC1Δ700 cDNA from pCL-CTC1-13xMyc, and then cloned into pCL-puro-FLAG-MYC plasmid using the In-Fusion® HD cloning kit (Takara). pBabe-puro-STN1-14xMyc was constructed by GenScript USA Inc. pCI-neo-His_6_-STN1-His_6_ plasmid was constructed by PCR amplifying STN1 cDNA from pCI-neo-Myc-STN1 using primers containing His_6_ sequences, and then cloning into the Xhol and Not1 sites of the pCI-neo vector. The pEAK8 Flag-CTC1, pEAK8 STN1, and pEAK8 TEN1-(His)_6_ expression plasmids were gifts from Dr. Liuh-Yow Chen (Institute of Molecular Biology, Academia Sinica (Chen et al, 2012)). For the co-expression of STN1 and TEN1, the cDNAs of STN1 and carboxyl-terminal (His)_6_-tagged TEN1 were cloned into pcDNA3.4 vector with a dual CMV promoter. The amino-terminal Flag-tagged CTC1Δ700N mutant was generated with deletion PCR mutagenesis by using pEAK8 Flag-CTC1 as the template. For the expression of human MRE11, the cDNA was amplified by PCR and then cloned into pcDNA 3.4 vector to harbor amino-terminal Flag-tagged and carboxyl-terminal (His)_6_-tagged MRE11. All constructs were sequenced to ensure sequence accuracy.

### Antibodies

| Antibodies | Source | Identifier |
| --- | --- | --- |
| Rat monoclonal anti-BrdU (reacting with CldU) | Abcam | Catalog No. ab6326;<br>RRID: AB_305426 |
| Mouse anti-BrdU (reacting with IdU) | BD Biosciences | Catalog No.347580;<br>RRID: AB_10015219 |
| Mouse anti-Myc (IF) | Millipore-sigma | Catalog No.05-419;<br>RRID :AB_309725 |
| Mouse monoclonal anti-Flag M2 | Millipore-sigma | Catalog No.F-1804-1mg;<br>RRID:AB_262044 |
| Rabbit Polyclonal anti-Flag | Millipore-sigma | Catalog No. F7425-2mg;<br>RRID :AB_439687 |
| Mouse monoclonal anti-c-Myc | Santa cruz<br>Biotechnology | Catalog No. sc-40;<br>RRID:AB_627268 |
| Rabbit Polyclonal anti-c-Myc | Bethyl | Catalog No. A190-105A;<br>RRID: AB_67390 |
| Rabbit Polyclonal anti-HA | Bethyl | Catalog No. A190-108A;<br>RRID: AB_67465 |
| Mouse monoclonal anti-RPA32 | Abcam | Catalog No. Ab2175;<br>RRID: AB_302873 |
| Rabbit Polyclonal anti-RPA32pS33 | Bethyl | Catalog No. A300-246A-T; RRID: AB_2180847 |
| Rabbit Polyclonal anti-ZRANB3 | Bethyl | Catalog No. A303-33A-T |
| Rabbit Polyclonal anti-SMARCAL1 | Bethyl | Catalog No. A301-616A-T; RRID: AB_2780017 |
| Rabbit Polyclonal anti-CTC1 | Abcam | Catalog No. ab175074 |
| Mouse monoclonal anti-TRF2 | Novus Biologicals | Catalog No. IMG-124A;<br>RRID: AB_317546 |
| Mouse polyclonal anti-STN1 | Abcam | Catalog No. ab89250;<br>RRID: AB_2156016 |
| Mouse monoclonal Anti-β-Actin | Millipore-sigma | Catalog No. A5441-100UL;<br>RRID: AB_476744 |
| Mouse monoclonal Anti-BRCA2 | Millipore-sigma | Catalog No. OP95-100UG<br>RRID: AB_213443 |
| Goat anti-Mouse Alexa Fluor 488 | ThermoFisher Scientific | Catalog No. A11029;<br>RRID: AB_138404 |
| Goat anti-Rabbit Alexa Fluor 550 | ThermoFisher Scientific | Catalog No.84541;<br>RRID: AB_10942173 |
| Goat anti-Rat Alexa Fluor 488 | ThermoFisher Scientific | Catalog No. A11006;<br>RRID: AB_2534074 |
| Goat anti-mouse Alexa Fluor 568 | ThermoFisher Scientific | Catalog No. A11031<br>RRID: AB_144696 |
| HRP Goat Anti-Rabbit | Vector Laboratory | Catalog No.PI-1000;<br>RRID: AB_2336198 |
| HRP Goat Anti-mouse | BD Pharmingen | Catalog No. 554002;<br>RRID: AB_395198 |

### Chemicals

| **Chemicals** | **Source** | **Identifier** |
| --- | --- | --- |
| X-tremeGENE™ HP DNA Transfection Reagent | Millipore Sigma | 6366244001 |
| Puromycin | VWR | AAJ61278 |
| Hygromycin B | Millipore Sigma | 400053 |
| Hydroxyurea | ThermoFisher Scientific | A10831 |
| Camptothecin | Millipore Sigma | 208925 |
| Methylmethanesulfonate | ThermoFisher Scientific | AC156890050 |
| Mirin | Millipore Sigma | M9948 |
| BrdU | MP Biomedicals | SKU 02100166-CF |
| IdU | Millipore Sigma | 54-42-2 |
| CIdU | MP Biomedicals | 105478 |
| EdU | Lumiprobe | 10540 |
| Biotin-azide | Jena Bioscience | CLK-1265-5 |
| Duolink in situ PLA probe anti-rabbit plus | Millipore Sigma | DUO92002 |
| Duolink in situ PLA probe anti-mouse minus | Millipore Sigma | DUO92002 |
| Duolink in situ detection reagents red | Millipore Sigma | DUO92008 |
| Mounting medium with DAPI | Vector Labs | H-1200 |
| Lipofectamine RNAiMax | ThermoFisher Scientific | Cat#13778030 |
| Supersignal West Pico | ThermoFisher Scientific | Cat#34078 |
| Supersignal West Femto | ThermoFisher Scientific | Cat#34095 |

### Experimental Models: Cell Lines

| **Cells** | **Source** | **Identifier** |
| --- | --- | --- |
| U2OS | ATCC | HTB-96; RRID: CVCL_0042 |
| HeLa | ATCC | CCL-2; RRID: CVCL_0030 |
| HEK293T | ATCC | CRL-3216; RRID: CVCL_0063 |
| BJ | ATCC | CRL-2522; RRID: CVCL_3653 |
| HCT116 | ATCC | CCL-247; RRID: CVCL_0291 |

### Software and Algorithms

| **Software** | **source** | **website** |
| --- | --- | --- |
| GraphPad Prism5 | GraphPad Software | https://www.graphpad.com/scientific-software/prism/; RRID: SCR_002798 |
| ImageJ | Department of Health and Human Services, NIDA | https://fiji.sc/ RRID: SCR_002285 |

### siRNA

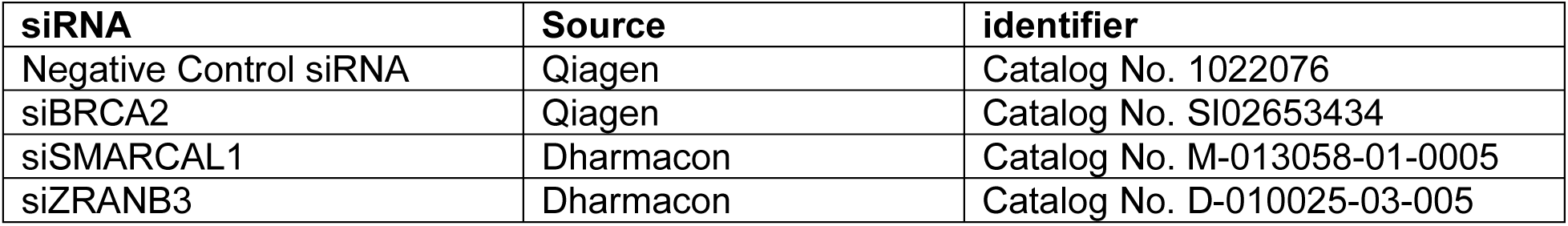

### shRNA

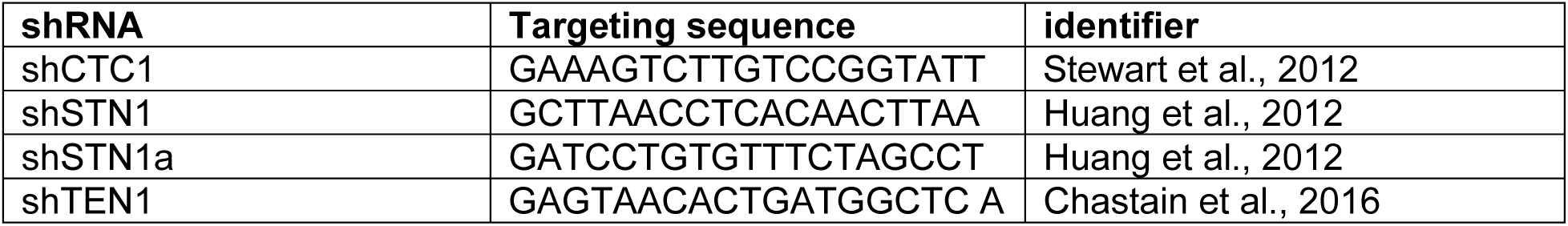

The above sequences were cloned into pSIREN-retro as previously described (Chastain et al, 2016).

### siRNA Transfections

siRNA used in this study are listed in the table above. All siRNAs oligos were transfected at a final concentration of 40 nM using Lipofectamine RNAiMAX (Invitrogen) according to the manufacturer’s protocol. Experiments were performed 60 hours post transfection.

### S-phase cell enrichment

Cells enrichment at mid-S phase was described previously (Chastain et al, 2016). Briefly, exponentially growing HeLa cells were treated with 2 mM thymidine for 14 h, followed by washing with pre-warmed medium for three times to remove thymidine. Cells were then cultured for an additional 16 h, and the second thymidine (2 mM) was added to medium for 12-16 h to arrest at the G1/S boundary. After washing with pre-warmed medium three times, cells were then released into fresh medium for 3 h to reach the mid-S phase. Cells were either collected immediately at mid-S phase, or treated with HU (2 mM) for an additional 3 h to induce fork stalling.

### Immunofluorescence (IF) staining

IF was carried out as described previously (Chastain et al, 2016), Briefly, cells were grown on cover slips or chamber slides, then either fixed directly with 4% paraformaldehyde (PFA) in PBS for 15 min, or permeabilized in 0.15% Triton X-100 for 2 min at room temperature prior to PFA fixation. Following fixation, cells were permeabilized with 0.15% Triton X-100 for 15 min, washed 3 times for 5 min with PBS, blocked with 10% BSA at 37 °C for 1 h in humidified chamber, then incubated with appropriate primary antibodies for overnight at 4 °C. Samples were then washed with PBS three times, incubated with secondary antibodies at room temperature for 1 h, and finally washed 3 times in PBS. Slides were then treated with cold ethanol series and dried in dark. Nuclei were visualized by counter staining with DAPI mounting medium (Vector Laboratories). Z-stack images were obtained at a 0.3 µm thickness per slice under Zeiss AxioImager M2 epifluorescence microscope with a 40x or 100x objective. Single Z-slice or max projection images were selected as representative images.

### ssDNA detection under the non-denaturing condition (BrdU staining)

To detect ssDNA under the non-denaturing condition, cells were cultured in media supplemented with 20 µM BrdU for 48 h, followed by 2 mM HU treatment for 6 h. After fixation with PFA, cells were permeabilized with 0.15% Triton X-100 for 15 min, washed 3 times for 5 min with PBS, blocked with 10% BSA at 37 °C for 1 h in humidified chamber, then incubated with anti-BrdU (Abcam) overnight at 4 °C. Regular IF procedures were then followed (see above).

### Detection of replicating cells (BrdU staining under the denaturing condition)

To detect replicating cells, cells were cultured in media supplemented with 10 µM BrdU for 1 h, and then rinsed with PBS two times. After fixation with 4% PFA for 15 min, cells were rinsed with PBS three times and then incubated with 1 M HCl at 37 °C for 20 min to denature DNA. Subsequently, denatured samples were washed with Na_3_PO_4_ (pH7.4) 3 times and then with PBS 5 times. Cells were then permeabilized with 0.15% Triton X-100 for 15 min. After washing with PBS three times, samples were blocked with 10% BSA at 37 °C for 1 h in humidified chamber and then incubated with anti-BrdU at 4 °C for overnight. Regular IF procedures were then followed (see above).

### Co-immunoprecipitation (Co-IP)

Cells were lysed in lysis buffer (0.1% NP-40, 50 mM Tris-HCl, pH 7.4, 50 mM NaCl, 2 mM DTT) supplemented with protease inhibitor cocktail (1 mM AEBSF, 0.3 µM aprotinin, 50 µM bestatin, 10 µM E-64, 10 µM leupeptin, 5 µM pepstain and 1 mM PMSF), sonicated on ice, and centrifuged (13,000 rpm, 15 min, 4 °C). The supernatants were precleared and immunoprecipitated with anti-Myc or anti-Flag antibody overnight at 4 °C with constant rotation. Beads were washed with cold lysis buffer 4 times, then resuspended in lysis buffer with SDS sample loading buffer, boiled for 10 min, and used immediately on SDS-PAGE for immunoblotting.

### SIRF assay

SIRF assay was performed as described previously (Roy et al, 2018). Briefly, Cells were exponentially grown on chamber slides and labelled with EdU for 8 min. Cells were washed with PBS once and fixed with 2% paraformaldehyde (PFA) for 15 min at room temperature. Slides were washed in Coplin jars 3 times for 5 min each with PBS. After permeabilization with 0.25% Triton X-100 in PBS for 15 min at room temperature, slides were washed 3 times for 5 min each with PBS, followed by incubation with click reaction cocktail (2 mM copper sulfate, 10 µM biotin-azide, and 100 mM sodium ascorbate) in humid chamber at 37 °C for 1 h. Slides were then washed 3 times for 5 min each with PBS and blocked with blocking buffer (10% BSA and 0.1% Triton X-100 in PBS) at 37 °C for 1 h. Primary antibodies were diluted in blocking buffer, dispensed onto slides, and incubated at 4 °C in humidified chamber. Slides were then washed with wash buffer A (0.01 M Tris, 0.15 M NaCl and 0.05% Tween-20, pH7.4) three times for 5 min each, and incubated with Duolink *in situ* PLA probe anti-mouse plus and anti-rabbit minus for 1 h at 37 °C. Following wash with wash buffer A three times for 5 min each, slides were incubated with Duolink ligation mix at 37 °C for 30 min, washed again with wash buffer A two times for 2 min each, incubated with Duolink amplification mix at 37 °C for 100 min, washed with wash buffer B (0.2 M Tris and 0.1 M NaCl) three times for 10 min each and one time in 0.01x diluted wash Buffer B for 1 min. Lastly, slides were dried in dark and nuclei were visualized by counter staining with DAPI mounting medium (Vector Laboratories). Images were obtained under Zeiss AxioImager M2 epifluorescence microscope with a 100x objective.

### DNA fiber analysis

DNA fiber assays were carried out as described previously (Nieminuszczy et al, 2016). Briefly, cells were first labeled with 25 µM CIdU for 30 min, followed by 250 µM IdU for 30 min, and then exposed to 4 mM HU for 5 h. Mirin (50 µM) was concomitantly added with HU as indicated. Subsequently, cells were harvested, resuspended in PBS at 1000 cells per µl, lysed in lysis buffer (200 mM Tris-HCl pH7.4, 50 mM EDTA, 0.5% SDS), and then DNA fibers were stretched on glass slides. Following fixation in methanol: acetic acid (3:1), slides were denatured with 2.5 M HCl for 80 min, washed with PBS and then blocked with 5% BSA in PBS (w/v) for 30 min. Nascent DNA labelled by CIdU and IdU were immunostained with anti-CldU and anti-IdU antibodies, then incubated with anti-rat Alexa Fluor 488 and anti-mouse Alexa Fluor 568 secondary antibodies (ThermoFisher Scientific). Coverslips were mounted using mounting medium (Vector Lab). Images were acquired with a Zeiss AxioImager M2 epifluorescence microscope at 40X magnification and analyzed using the Image J software. In all DNA fiber experiments in this study, at least two independent treatments and fiber experiments were performed to ensure reproducibility. Results from one set of experiment are shown in each graph. Scatter plots indicates IdU/CIdU tract length ratios for individual replication forks.

### Cell survival measurement

Eight hundred cells were seeded in 6-well plates in triplicate one day prior to treatment. Cells were then treated with indicated concentrations of CPT, and treatment was repeated every 4 days at 37 °C. For MMS treatment, cells were incubated with the indicated concentrations of MMS for 48 h, then MMS was removed from media. After 10 days of incubation, the medium was removed, cells were washed with PBS, then fixed and stained with the crystal violet solution (0.1% crystal violet, 1% methanol and 1% formaldehyde). Cell viability was calculated by normalizing the colony numbers of treated samples to untreated samples. For each drug treatment, three independent experiments were performed.

### Expression and purification of the human CST complex and human MRE11

#### Expression

pEAK8-CTC1 WT or pEAK8-CTC1Δ700N and pcDNA3.4-STN-TEN1-(His)_6_ plasmids were co-transfected into Expi293F cells according to instruction manual from ExpiFectamine 293 kit (ThermosFisher Scientific). Cells were harvested by centrifugation after 72 h transfection.

#### Purification

All protein purification steps were carried out at 4°C. Cells were suspended in buffer A (25 mM Tris-HCl, pH 7.5, 10% glycerol, 0.01% Igepal, 150 mM KCl and 1 mM 2-mercaptoethanol) supplemented with protease inhibitors (1 mM PMSF and Benzamidine, 3 μg/ml each of aprotinin, chymostatin, leupeptin and pepstatin A). Cells were lysed by sonication, followed by centrifugation at 40,000x g for 1 h. Lysates were supplemented with 10 mM imidazole and incubated with Ni^2+^ NTA-agarose resin (QIAGEN) for 3 h. The resin was then poured into an open column and washed with buffer A containing 10 mM imidazole twice. CST complex was eluted with buffer A (without 2-mercaptoethanol) containing 200 mM imidazole. The eluate was then loaded into the anti-Flag M2 affinity gel packaged column (Sigma) and washed with buffer A (without 2-mercaptoethanol) twice. The CST complex was eluted with buffer A (without 2-mercaptoethanol) containing 3x Flag peptide (100 µg/ml). The affinity-purified CST complex was then loaded on Superdex 200 increase 10/300 GL column (GE Healthcare) equilibrated with butter A. The peak fractions were pooled and concentrated, then divided into small aliquots and stored at −80°C. The CTC1 Δ700N complex was purified using the same protocol.

For the human MRE11 purification, transient transfection of Flag-MRE11-(His)_6_ plasmid was also performed in Expi293F cells for 72 h. Cells were harvested and suspended with buffer A containing 500 mM KCl and the protease inhibitors as described above. After sonication and centrifugation, lysates were supplemented with 10 mM imidazole and incubated with Ni^2+^ NTA-agarose resin for 3 h. The resin was then poured into an open column and washed with buffer A containing 500 mM KCl and 10 mM imidazole twice. MRE11 was eluted with buffer A (without 2-mercaptoethanol) containing 200 mM imidazole. The eluate was then loaded into the anti-Flag M2 affinity gel packaged column (Sigma) and washed with buffer A (without 2-mercaptoethanol) twice. The MRE11 protein was eluted with buffer A (without 2-mercaptoethanol) containing 3x Flag peptide (100 µg/ml), and then the eluate was fractionated in a 1 ml MonoQ column (GE Healthcare), using a 30 ml gradient of 150-1000 mM KCl in buffer A. The peak fractions containing pure MRE11 were pooled and concentrated, then divided into small aliquots and stored at −80°C.

#### DNA substrates

Fluorescence-labeled 5’ overhang DNA substrate was prepared by annealing the synthetic oligonucleotides described below. Oligo 1 (60 nt) with the modified phosphorothioate bond (labeled with an asterisk) at both ends to prevent the non-specific nucleases digestion was purchased from IDT: 5’-A*C*G*C*T*GCCGAATTCTACCAGTGCCTTGCTAGGACATCTTTGCCCACCTGCAGGTTC*A* C*C*C*-3’ Oligo 2 (25 nt) with modified Cy3 fluorescence dye at the 5’ end was purchase from Genomics: 5’-Cy3-GGGTGAACCTGCAGGTGGGCAAAGA-3’

Briefly, equal amounts of oligonucleotides were mixed in the annealing buffer (50 mM Tris pH 7.5, 10 mM MgCl_2_, 100 mM NaCl, and 1 mM DTT) and heated at 80 °C for 3 min. The mixed reaction was subsequently transferred to 65°C for 30 min and cooled down slowly to room temperature. The annealed substrate was purified from a 10% native polyacrylamide gel by electro-elution and filter-dialyzed in an Amicon ultra-4 concentrator (Millipore, NMWL 10 kDa) at 4°C into TE buffer (10 mM Tris-HCl, pH 8.0, and 0.5 mM EDTA). The substrate concentration was quantified by using absorbance at 260 nm and the molar extinction coefficients of the substrate with Cy3 calculated by Molbiotools online software.

### Electrophoretic mobility shift assay

The fluorescence-labeled 5’ overhang DNA substrate (80 nM) was incubated with indicated amounts of CST complex (CTC1 wild-type or CTC1 Δ700N) in 10 µl buffer B (35 mM Tris-HCl, pH 7.5, 1 mM DTT, 100 ng/µl BSA, and 50 mM KCl) at 37°C for 5 min. The reaction mixtures were then electrophoresed on 6% native TBE-PAGE with 1× TBE buffer (89 mM Tris, 89 mM borate, and 2 mM EDTA, pH 8) at 100 V for 30 min at 4°C. Gels were analyzed by BioSpectrum 810 imaging system with a 533-587 nm filter (UVP).

### MRE11 degradation assay

Fluorescence-labeled 5’ overhang DNA substrate (80 nM) was incubated with indicated amounts of CST complex (CTC1 wild-type or CTC1 Δ700N) in 10 μl buffer B containing 2.5 mM MgCl_2_, 1 mM ATP, and 1 mM MnCl_2_ at 37°C for 5 min, followed by incubation of purified human MRE11 (200 nM) at 37°C for 20 min. Reactions were stopped by incubation with 2.5 µl stop buffer (50 mM EDTA, 0.4% SDS and 3.2 mg/ml proteinase K) at 37°C for 15 min. Samples were then mixed with the equal volume 2× denature dye (95% formamide, 0.1% Orange G, 10 mM Tris-HCl, pH 7.5, 1 mM EDTA, and 12% Ficoll PM400), heat denatured at 95°C for 10 min, and analyzed on 27% denature TBE-Urea-PAGE (7M Urea) with 1× TBE buffer at 300 V for 50 min at 55°C. Gels were analyzed by BioSpectrum 810 imaging system with a 533-587 nm filter. The intensity of DNA was quantified by the Image J software. Images shown represent results of independent experiments repeated at least three times. Data were analyzed and presented with the PRISM 7 software and shown as mean ± S.D.

### Prognostic value analysis

Kaplan–Meier analysis was performed using KM-plotter online software (Gyorffy et al, 2010) (http://kmplot.com/analysis/). The relationship of gene expression and relapse free survival (RFS) (n = 3.955) was evaluated in an integrated multi-study breast cancer microarray data set containing 13 breast cancer expression profiling data sets from GEO. Kaplan–Meier estimates of RFS (relapse-free survival) were calculated by setting the software to look for the optimal cut-off for separation of patients into the higher- and lower-tertile expressing groups. The hazard ratio, log-rank P value, and number of patients in each group are shown on the KM plot for each gene.

## Acknowledgement

We thank Liuh-Yow Chen (Academia Sinica, Taiwan) for constructs. The work is supported by NIH R01GM112864 and R01CA234266 to W.C., and Academia Sinica, National Taiwan University, and Taiwan Ministry of Science Technology (MOST 108-2321-B-002-054) to P.C.

## Author contributions

X.L. performed the majority of experiments except for the in vitro studies. K-H.L. purified CST and MRE11 and carried out the in vitro experiments. O.S. participated in various experiments and collected data. M.C. performed IF experiments, generated K-M plots, and collected data. W.C. and P.C. conceived and directed the study. X.L., K-H.L., P.C., W.C. wrote the paper.

